# Endogenous opioid receptor system mediates costly altruism in the human brain

**DOI:** 10.1101/2023.12.06.569026

**Authors:** Jinglu Chen, Vesa Putkinen, Kerttu Seppälä, Jussi Hirvonen, Kalliopi Ioumpa, Valeria Gazzola, Christian Keysers, Lauri Nummenmaa

**Author notes:** Address for correspondence Jinglu Chen, Turku PET Centre c/o Turku University Hospital FI-20520 Turku, Finland.

## Abstract

Functional neuroimaging studies suggest that a large-scale brain network transforms others’ pain into its vicarious representation in the observer, potentially modulating helping behaviour. The neuromolecular basis of individual differences in vicarious pain and helping, however, have remain poorly understood. Here we investigated the role of the endogenous μ-opioid receptor (MOR) system – known for its role in analgesia and sociability – in altruistic costly helping. MOR density was measured with high-affinity agonist radioligand [^11^C]carfentanil. In a separate fMRI experiment, participants could choose to donate money to reduce the pain of the confederate who was subjected to electric shocks of varying intensity. We found that subjects were in general willing to engage in costly helping, and haemodynamic activity in amydala, anterior insula, anterior cingulate cortex, striatum, primary motor cortex, primary somatosensory cortex, and thalamus increased when participants witnessed the pain of others. These haemodynamic responses were negatively associated with MORs availability in the striatolimbic and cortical emotion circuits. In turn, haemodynamic responses during helping were positively associated with MOR availability in the anterior cingulate cortex and hippocampus. Altogether these data suggest that the endogenous MOR system modulates the processing of altruistic behaviour and costly helping in the human brain.

## Introduction

Prosocial behavior is prevalent in humans and animals. Social animals share emotions with each other, comfort peer’s distress, and help others in times of need (De Waal & Preston, 2017; Keysers et al., 2022). Humans help each other in daily life even without genetic relatedness or obvious direct profit (Batson, 2010, 2011; Batson et al., 1997). Such altruistic behavior by definition benefits the receiver, but also has a range of positive outcomes for the helper, including improved academic performance and social preferences (Caprara et al., 2000), increased acceptance by peers at school (Deković & Gerris, 1994), and higher life satisfaction (Caprara & Steca, 2005). The evolutionary routes of altruism are a topic of enduring interest (Dovidio & Penner, 2003). For more information, see review Laguna et al., 2020. A prominent hypothesis for the proximate causes for altruistic helping holds that if we empathically activate our own pain when witnessing the pain of others, mechanisms that have evolved to motivate us to prevent damage to ourselves will also motivate us to prevent pain and damage to others. Accordingly, empathy can be hypothesized to drive prosociality (Batson et al., 2014; Decety et al., 2016; Keysers & Gazzola, 2014; Smith, 1759).

The neurocognitive link between empathy and prosociality has been investigated by quantifying responses in brain regions associated with empathy while witnessing the pain of others, and correlating these measures with individual differences in helping. Studies leveraging a range of different methods have consistently shown that seeing or hearing others in distress compared to neutral states engages a core network including the anterior insula (aIns) and adjacent frontal operculum, the mid- to anterior-cingulate cortex, amygdala and, when the somatic source of the pain is salient, the primary and secondary somatosensory cortices (SI and SII) (Bufalari et al., 2007; De Waal & Preston, 2017; Keysers et al., 2010; Krishnan et al., 2016; Lamm et al., 2011; Paradiso et al., 2021). Based on the details of the task and the nature of how the pain is witnessed, this extends into a larger network also encompassing primary motor cortex (M1), extensive areas in frontal lobe and parietal lobe, dorsolateral prefrontal cortex (dlPFC), ventromedial frontal cortex (vmPFC), fusiform gyrus, temporal pole, precuneus, thalamus, caudate, and putamen(Jauniaux et al., 2019; Timmers et al., 2018). Particularly the ACC and aIns are activated both when subjects experience first-hand pain and when they see others experiencing pain, suggesting that these regions underlie the transformation of others’ pain from visual and auditory inputs into sensorimotor formats (Bufalari et al., 2007; Jackson et al., 2005, 2006; Krishnan et al., 2016; Lamm et al., 2011; Saarela et al., 2007). The ACC has been shown to contain single neurons that respond to both witnessing and experiencing pain in rodents(Carrillo et al., 2019).

To investigate whether this recruitment of regions involved in the witnesses’ own pain plays a role in motivating helping, some studies have gone beyond simply measuring brain activity while passively witnessing the pain of others and provided the participant with an opportunity to help. One study offered participants the opportunity to relieve other people’s pain by taking some of that pain onto themselves and found that anterior insular activity predicted the willingness to help ingroup members and nucleus accumbens activity predicted reluctance to help outgroup members (Hein et al., 2010). Another found that reduced functional connectivity between the insula and ACC characterized participants that decide to help a person in need in a virtual reality scenario (Zanon et al., 2014). Activity in somatosensory and insular region while witnessing pain predicted later charitable donations, albeit in ways that depended on socioeconomic status (Ma et al., 2011). An approach that has been particularly successful in identifying brain regions with activity quantitatively associated with helping, involves offering participants a certain financial endowment, and enabling them to forfeit some of that endowment on a trial-by-trial basis to reduce the pain of a victim in an immersive experimental setting (FeldmanHall et al., 2012). These studies have found a variety of nodes also involved with self-pain to relate to individual or trial-by-trial differences in helping, including the dorsolateral prefrontal cortex, orbital frontal cortex (FeldmanHall et al., 2015), SI (Gallo et al., 2018), vmPFC (FeldmanHall et al., 2013; Fornari et al., 2023), insula, ACC, and amygdala (Caspar et al., 2022).

Although these financially costly helping paradigms thus start to provide some traction on which neural networks have BOLD activity that correlate with costly helping, the molecular basis of empathy and costly helping, and their individual differences, remain poorly understood. Multiple lines of evidence point towards the critical role of the endogenous opioid system and particularly the endogenous 𝜇-opioid receptor (MOR) system in the first-person experience of pain and in differences in empathy and prosociality. Among the three classes of opioid receptors (μ, δ, and κ), the μ receptors mediate the effects of endogenous β-endorphins, endomorphins, enkephalins, and various exogenous opioid agonists(Henriksen & Willoch, 2008). The predominant action of µ-opioids in the central nervous system is inhibitory, but they can also exert excitatory effects, and MORs are expressed widely throughout the human emotion circuits(Kantonen, Karjalainen, Isoj, et al., 2020; Nummenmaa & Tuominen, 2018). Opioids are well known for their role in antinociception, and genetic deletions of the MOR in mice abolishes the analgesic effect of opioid agonists (Darcq & Kieffer, 2018), demonstrating that this receptor is the sole pathway for the analgesic effects of these drugs. The effect of opioids however goes beyond nociception: opioid agonists decrease and antagonists increase social motivation in macaques (Fabre-Nys et al., 1982; Keverne et al., 1989), and in humans, the opioid system modulates both positive and negative emotions (Nummenmaa et al., 2020; Nummenmaa & Tuominen, 2018)

With regard to empathy, recent experiments have shown that activating the MOR system using placebo analgesia reduces first-hand pain ratings, how much pain participants perceive in others, and how unpleasant they find witnessing such pain (Rütgen, Seidel, Riečanský, et al., 2015; Rütgen, Seidel, Silani, et al., 2015), and this effect can be blocked using a MOR antagonist (Rütgen, Seidel, Silani, et al., 2015). Also, the long term use of opioids, known to dysregulate the MOR system, leads to reduced pain ratings in others (Kroll et al., 2021). One study also showed that placebo analgesia reduces helping (Hartmann et al., 2022). Positron emission tomography (PET) with [^11^C] carfentanil, a synthetic, highly specific MOR agonist allows in vivo imaging of opioid receptors in humans. Studies with PET have demonstrated endogenous opioid release in the thalamus following acute pain in a dose-dependent manner (Bencherif et al., 2002). Individual differences in MOR availability also link with pain sensitivity: participants with lower MOR availability have a higher sensitivity to pain (Hagelberg et al., 2012). Finally, and critically with respect to the present study, PET studies have linked opioid receptor availability with vicarious pain perception and sociability. First, MOR availability is negatively associated with haemodynamic responses to seeing others in pain (Karjalainen et al., 2017). The MOR system is also activated during positive social interactions such as laughing together (Manninen et al., 2017), and MOR availability is positively correlated with prosocial motivation as indexed by social attachment styles (Nummenmaa et al., 2015; Turtonen et al., 2021). Against this background, it could be hypothesized that the MOR system would be a crucial molecular pathway for altruistic, costly helping, but currently this hypothesis lacks direct in vivo evidence.

### The current study

Here we investigated whether individual differences in the MOR availability at rest translate into measurable differences in the willingness to forfeit money to reduce pain to others, using the established “costly” helping paradigm of Gallo et al.. In the fMRI experiment, participants could choose to donate money to reduce the pain of the confederate who was subjected to electric shocks of varying intensity. In a separate scanning session, the participants underwent a baseline PET scan with the MOR specific radioligand [^11^C]carfentanil. We found that people were in general willing to engage in costly helping; The fMRI results revealed that activity in amydala, aIns, anterior cingulate cortex, striatum, primary motor cortex, primary somatosensory cortex, and thalamus increased when participants saw the confederate in pain. These haemodynamic responses had amplitudes that differed across individuals in ways that correlated with the availability of MORs in the striatolimbic and cortical emotion circuits. Altogether these data suggest that endogenous MOR system contributes to altruistic behavior and its individual differences.

## Materials and methods

### Participants

Thirty healthy Finnish women (mean ± SD age: 24.7±5.65 years, range 19-42) with normal or corrected to normal vision were recruited in the study. To maximize statistical power of the study, only females were included because the MOR system shows a high level of sexual dimorphism (Kantonen, Karjalainen, Isoj, et al., 2020). In addition, research on sex differences in empathy and pain shows that women are better at judging emotional signals (Hall & Matsumoto, 2004), show higher emotional responsivity (Christov-Moore et al., 2014), evaluate others’ pain as more intense (Robinson & Wise, 2003), are more empathic than men (Hoffman, 1977), and are not influenced by trait harm sensitivity (FeldmanHall et al., 2016). All subjects participated in the fMRI scan and 14 of them participated in the PET scan. PET and MRI scans were conducted on separate days. Exclusion criteria included medications affecting the central nervous system, mood or anxiety disorders, psychotic disorders or neurological conditions, substance abuse and standard MRI and PET exclusion criteria. Structural brain abnormalities that are clinically relevant or could bias the analyses were excluded by a consultant neuroradiologist. The study was approved by the ethics board of the hospital district of Southwest Finland and conducted according to Good Clinical Practice and the Declaration of Helsinki (Approval number: 51/1801/2019). All subjects signed written informed consent and were informed that they had the right to withdraw at any time without giving a reason. The participants were compensated for their time and travel costs.

### Experimental design and stimuli for fMRI

The fMRI study was run using the costly helping paradigm of Gallo et al. (2018). The participants were led to believe that they were witnessing the distress of another person in real time. Before the experiment, participants met the confederate and were explained that the experiment would be performed in two separate rooms connected by a video camera. They were invited to draw lots to determine who would undergo the fMRI measurement while seeing the other participant receiving the electric shocks. The confederate was always chosen to receive the electric shocks. During the scan, participants, believing to witness the pain of the victim through a closed-circuit television, actually they viewed pre-recorded videos of the confederate receiving the painful simulation. All participants were debriefed at the end of the experiment.

The task was run by Presentation software (https://www.neurobs.com/) and the trial structure of the experiment is shown in **Figure 1**. On each trial, subjects first saw a video of the confederate receiving a painful electric shock (1^st^ video). The intensity of expressed pain ranged randomly from 2 (mild pain) to 6 (moderate pain) out of ten. After the video, subjects could decide how much money they were willing to donate on that trial to reduce the intensity of the second shock in that trial, to do that, one button was held by the participant in each hand, one representing increasing money and the other representing decreasing money. For each trial, they were given 6€. They knew that if they did not donate any money, the second shock would have the same intensity as the first, whilst, for every donated 1€, the second shock would be reduced by one point on the 10-point pain scale. Participants were told that they can keep the undonated money of all trials divided by 10 as their extra compensation after the experiment. The subjects then saw the 2^nd^ video of the confederate receiving the second electric shock whose intensity reflected the first shock minus the donation (see supplementary for more information on videos). There were two imaging runs with 15 trials in each.

**Figure 1.**
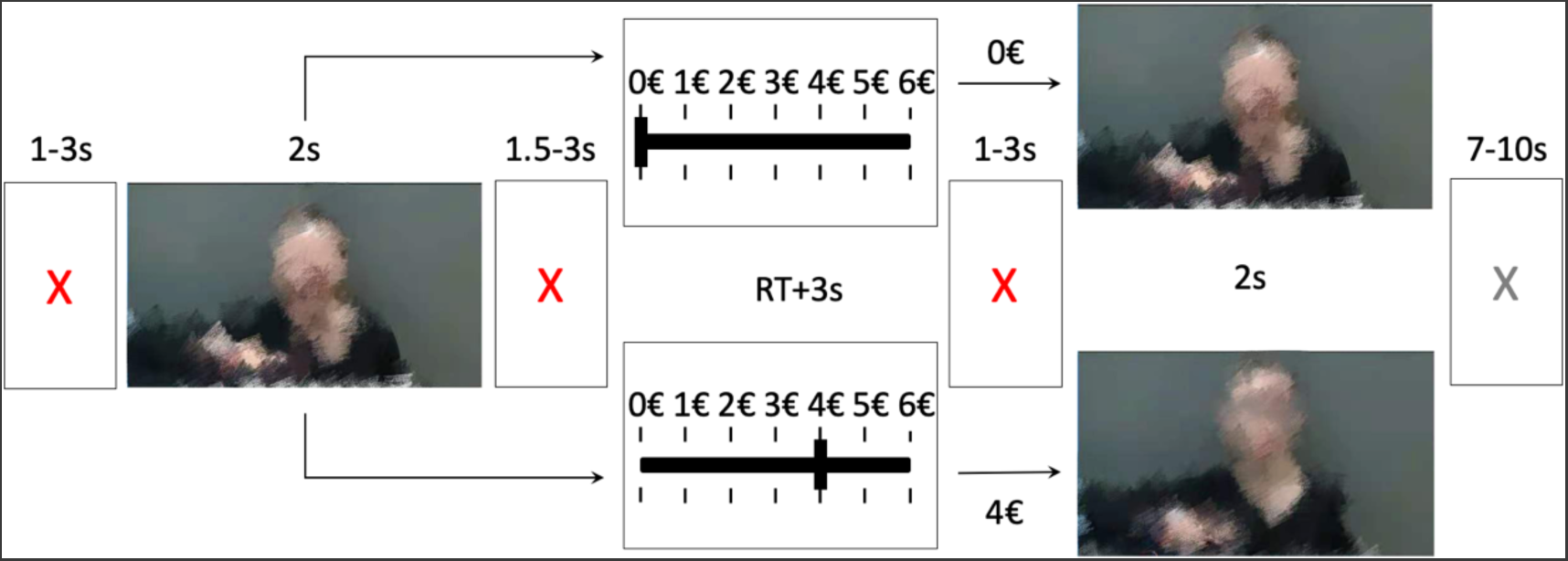
Trial structure. A red fixation cross was shown for 1-3s and was followed by 1^st^ video for 2s. Then another red fixation cross was shown for 1.5-3s. The donation phase was self-paced and was followed by a red fixation cross (1-3 s). Next, the post-donation video was shown for 2s followed by a gray fixation cross (7-10s). The snapshots from the videos shown on the right illustrate two possible scenarios from the 6 possibilities.

### MRI data acquisition and analysis

fMRI data were acquired with the 3T scanner (SIGNA, Premier, GE Healthcare, Waukesha, WI, USA) at the University Hospital of Turku. T2*-weighted functional images were collected with echo-planner imaging (EPI) sequence (45 slices; slice thickness = 3 mm; TR = 2600 ms; TE = 30 ms; flip angle =75°; FOV = 24 mm; voxel size = 3×3×3 mm^3^). T1-weighted structural images were collected with voxel size of 1×1×1 mm^3^. MRI data were preprocessed with fMRIPrep 1.3.0.post2 (Esteban et al., 2019). The following preprocessing was performed on the T1-weighted (T1w) image: correction for intensity, skull-stripping, brain surfaces reconstruction, refined brain mask estimating, cortical gray-matter segmentations, spatial normalization to the ICBM 152 Nonlinear Asymmetrical template version 2009c using nonlinear registration with antsRegistration (ANTs 2.2.0), and brain tissue segmentation. The following preprocessing was performed on the functional data: generation of reference volume and its skull-stripped version, co-registration to the T1w reference, slice-time correction, spatial smoothing with an isotropic, Gaussian Kernel of 6mm FWHM (full-width half-maximum), automatic removal of motion artifacts using ICA-AROMA (Pruim et al., 2015), and resampling to MNI152NLin2009cAsym standard space, and principal components are estimated after high-pass filtering the preprocessed BOLD time-series for the two CompCor variants: temporal and anatomical.

fMRI data were analyzed in SPM12 (Wellcome Trust Center for Imaging, London, UK, https://www.fil.ion.ucl.ac.uk/spm/). To investigate regions activated by i) seeing pain in the 1^st^ video, ii) donating, and iii) seeing pain in the 2^nd^ video, first-level general linear models (GLM) were estimated by modeling the 1^st^ video, donation phase, and the 2^nd^ video by using boxcar regressors in the design matrix. Donation size (trial-wise donations for each subject) was entered as parametric modulator for the 1^st^ video. Subject-wise contrast images were then generated for main effects of 1^st^ video, donation phase, 2^nd^ video. Additionally, a subtraction contrast was computed for 1^st^ vs. 2^nd^ video and the parametric effect of donation size for 1^st^ video. The contrast images were then subjected to second-level (random effects) analysis. Results are shown after FWE correction for cluster-size, by initially thresholding statistical maps at *p_unc_*<0.001, identifying the FWEc minimum cluster-size value for FWE correction at the cluster-size level, and then thresholding the statistical maps again at *p_unc_*<0.001 and k=FWEc-1.

### PET data acquisition and analysis

PET data were acquired during resting baseline with a GE Discovery Molecular Insights DMI PET/CT, GE Healthcare, Waukesha WI in Turku PET Center. The high-affinity agonist radioligand [^11^C]carfentanil was used to measure brain μ-opioid receptor availability. After intravenous radioligand injection (250.6±10.9 MBq), radioactivity in the brain was measured by the PET scanner for 51 minutes with increasing frame length (3×60s, 4×180s, 6×360s) using in-plane resolution of 3.77mm FWHM (Full Width Half Maximum) and tangential 4.00mm FWHM. All subjects lay supine in the PET scanner throughout the study. Data were corrected for dead-time, decay, and measured photon attenuation. In-house MAGIA pipeline was used to preprocess PET images (Kantonen, Karjalainen, Isojärvi, et al., 2020; Karjalainen et al., 2020).

Radiotracer binding was quantified using binding potential (*BP*_ND_), calculated as the ratio of specific binding to non-displaceable binding in the tissue (Innis et al., 2007). This outcome measure is not confounded by differences in peripheral distribution or radiotracer metabolism, or alterations in brain perfusion. *BP*_ND_ is traditionally interpreted by target molecule density (*B*_max_), although [^11^C]carfentanil is also sensitive to endogenous neurotransmitter release. Accordingly, the *BP*_ND_ for the tracer should be interpreted as the density of the receptors unoccupied by endogenous ligand (i.e., receptor availability). Binding potential was calculated by applying the basis function method (Gunn et al., 1997) for each voxel using the simplified reference tissue model (Lammertsma & Hume, 1996), with occipital cortex serving as the reference region (Frost et al., 1989). The parametric images were spatially normalized to MNI-space via segmentation and normalization of T1-weighted anatomical images, and finally smoothed with an 8-mm FWHM Gaussian kernel. PET imaging with [^11^C]carfentanil has high test-retest stability (Hirvonen et al., 2009). PET imaging always preceded fMRI to avoid potential impact of the fMRI tasks on measured MOR levels (mean ± SD day: 84±62 days, range 4-190).

### ROI selection

The average tracer *BP*_ND_ was quantified in 17 anatomical *a priori* regions of interest (ROI) involved in vicarious pain and empathy: amygdala, caudate, cerebellum, dorsal anterior cingulate cortex, inferior temporal gyrus, insula, middle temporal gyrus, nucleus accumbens, orbitofrontal cortex, pars opercularis, posterior cingulate cortex, putamen, rostral anterior cingulate cortex, superior frontal gyrus, superior temporal gyrus, temporal pole, and thalamus. The selection was based on previous studies on the effects of MORs on vicarious pain and arousal (Karjalainen et al., 2017, 2019). The ROIs were derived separately for each subject from the T1-weighted MR images using FreeSurfer (http://surfer.nmr.mgh.harvard.edu/).

### PET-fMRI Fusion Analysis

Three approaches were taken to characterize the interactions between MOR availability and BOLD responses in pain perception and costly altruism. In the first two approaches, a principal component analysis (PCA) was used to reduce the dimensionality in the *BP*_ND_ values across our ROIs. This was done because regional MOR availability has high autocorrelation (Tuominen et al., 2014), thus PCA would increase the power of our analyses by reducing the multiple comparison correction that would otherwise reduce power. We found the first 3 PCs to explain >90% of the variance, with 61%, 22% and 7% of variance explained, respectively. To identify voxels with BOLD responses that depend on individual differences in the overall MOR signal, we used the first PC to predict the voxel-wise BOLD responses to the 1^st^ video with donation size as a parametric modulator and the donation phase, separately. Specifically, we used the same parametric model as the fMRI analysis in the first-level model, and input the first PC in the second-level model for 1^st^ video and donation, in separate models. Second, to explore the relationship between individual MOR differences and responses in the vicarious pain observation network, we used the affective vicarious pain signature (AVPS) to dot-multiply the 1^st^ level beta maps to 1^st^ video with donation size as a parametric regressor for each participant (Zhou et al., 2020), thereby reducing each parameter estimate volume to a scalar value, and computed the correlation between the resulting value and the first 3 MOR PCs. Finally, to replicate previous studies on the links between MOR availability and haemodynamic responses to vicarious pain and arousal (Karjalainen et al., 2017, 2019), the voxel-wise BOLD responses to donation size in 1^st^ video were predicted with ROI-wise [^11^C]carfentanil binding potentials using whole-brain linear regression analysis with a statistical threshold set at *p* < 0.05, FWE-corrected at cluster-level. We then computed a cumulative map of the binarized MOR × BOLD beta maps to highlight regions whose BOLD responses were most consistently dependent on regional MOR availability.

## Results

### Behavioral results

Participants donated money in all intensity conditions (M = 2.826, SD = 1.964), and a linear mixed model confirmed that donations increased as a function of the shock intensity shown in the first video, *β* = 0.877, *SE* = 0.023, *t* = 37.591, *p* < 0.001, intercept is 0.198 (**Fig. 2**). The slope (beta=0.877) indicates that participants overall donate money to reduce shock intensity by 88%, adapting their donations very precisely to how much pain was at stake. In addition, in the linear mixed model, we added subject as a random effect. The variance of the effect from subject is 1.043, SD is 1.021.

**Figure 2.**
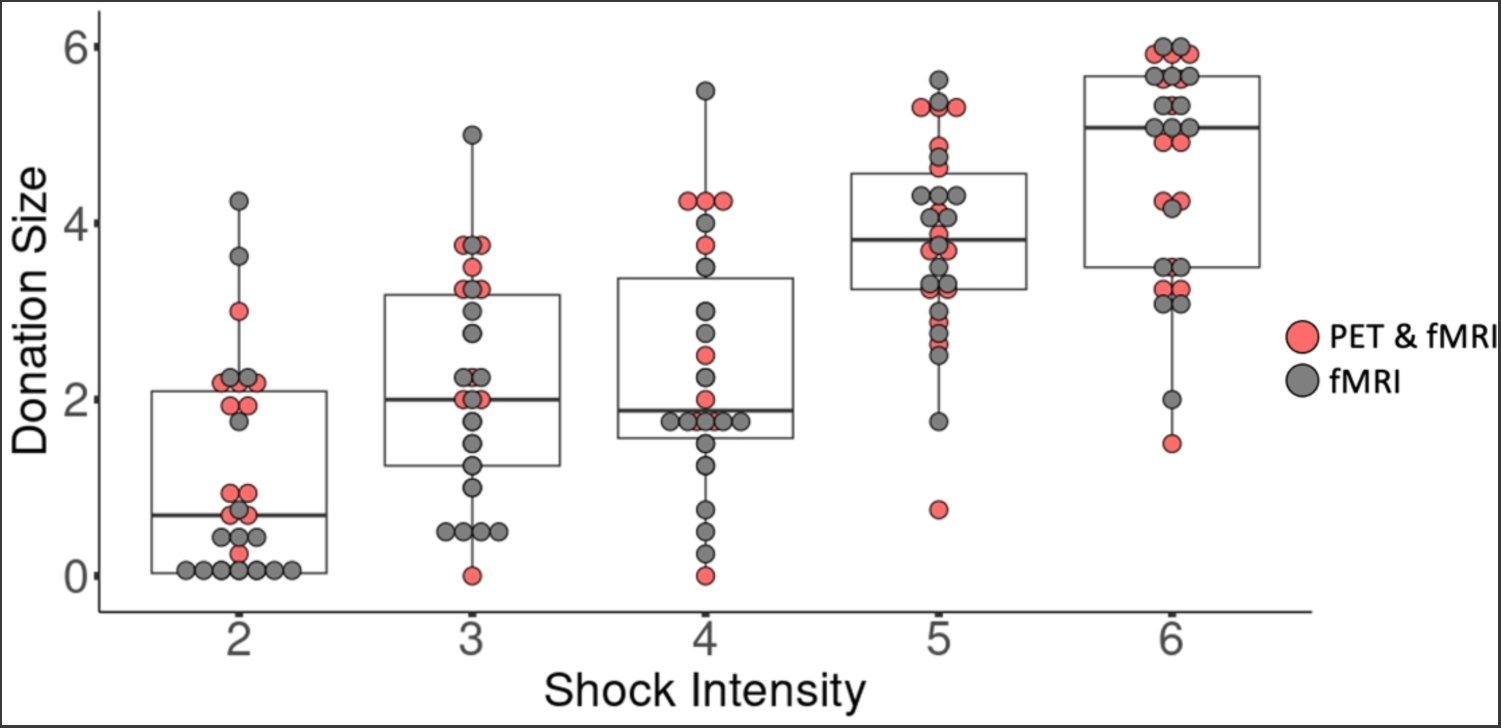
Donation size increased with increasing shock intensity. Each dot represents the average donation of a participant for all the stimuli of that intensity. N=30 participants. The box represents the 1st, 2nd and 3rd quartile, the whiskers the highest and lowest value within 1.5 interquartile range. Red dots represent participants who undertook both PET and fMRI scans. Gray dots represent participants with only fMRI scans.

To explore whether donations depended on the MOR availability for the 14 participants who also underwent the PET scan, for each participant, we calculated the slope and intercept linking the participants donation with the intensity of shocks shown in 1^st^ video (donation=slope*intensity+intercept). We then explored whether individual differences in slope or intercept were associated with individual differences in the MOR availability using the first 3 principal components of the MOR. We first did so using a multiple regression including all three PCs, which did not yield significant results for the slope (F(3,10)=0.036, p=0.990) or intercept (F(3,10)=0.552, p=0.658). Performing Bayesian correlation tests between the slope and intercept and each individual PCA (**Table 1**), confirms that this data is more likely under null hypothesis of no association. Together, this suggest that the actual donations do not depend on MOR availability in our ROIs.

**Table 1.** Lack of association between individual differences in slope or intercept and MOR PCs. For each of the 14 PET participants, we estimated the slope and intercept of the regression donation=slope*intensity+intercept. We then calculated the Pearson’s r value between the 14 slope (top) or intercept (bottom rows) with Jasp (https://jasp-stats.org/).

| donation | MOR | Pearson's r | p | BF<sub>10</sub> |
| --- | --- | --- | --- | --- |
| slope | pca1 | 0.075 | 0.800 | 0.339 |
| slope | pca2 | 0.013 | 0.965 | 0.329 |
| slope | pca3 | -0.070 | 0.813 | 0.338 |
| intercept | pca1 | -0.100 | 0.734 | 0.347 |
| intercept | pca2 | -0.136 | 0.642 | 0.363 |
| intercept | pca3 | 0.337 | 0.239 | 0.621 |

We therefore explored what brain circuits are involved in making the donation decisions, and whether these circuits vary based on MOR availability even if the outcomes of the decisions do not.

### BOLD-fMRI responses to vicarious pain perception

Full random effects results maps are available on NeuroVault (https://identifiers.org/neurovault.collection:14151). Consistent with previous studies, whole-brain general linear model analysis (GLM) revealed that anterior insula, anterior cingulate cortex (ACC) and thalamus were activated when seeing the confederate in pain in the 1^st^ video (**Fig. 3**). Additional activations were observed in the amygdala and striatum, which are key nodes of the emotion and reward networks. We next modelled the BOLD signal with the donation size in the 1^st^ video as a regressor. Consistent with previous research, insula, ACC activated more strongly as the donation increased (**Fig. 4**) (Ioumpa et al., 2023; Jackson et al., 2005, 2006; Karjalainen et al., 2017; Lamm et al., 2011; Saarela et al., 2007; Singer et al., 2004; Soyman et al., 2022).

**Figure 3.**
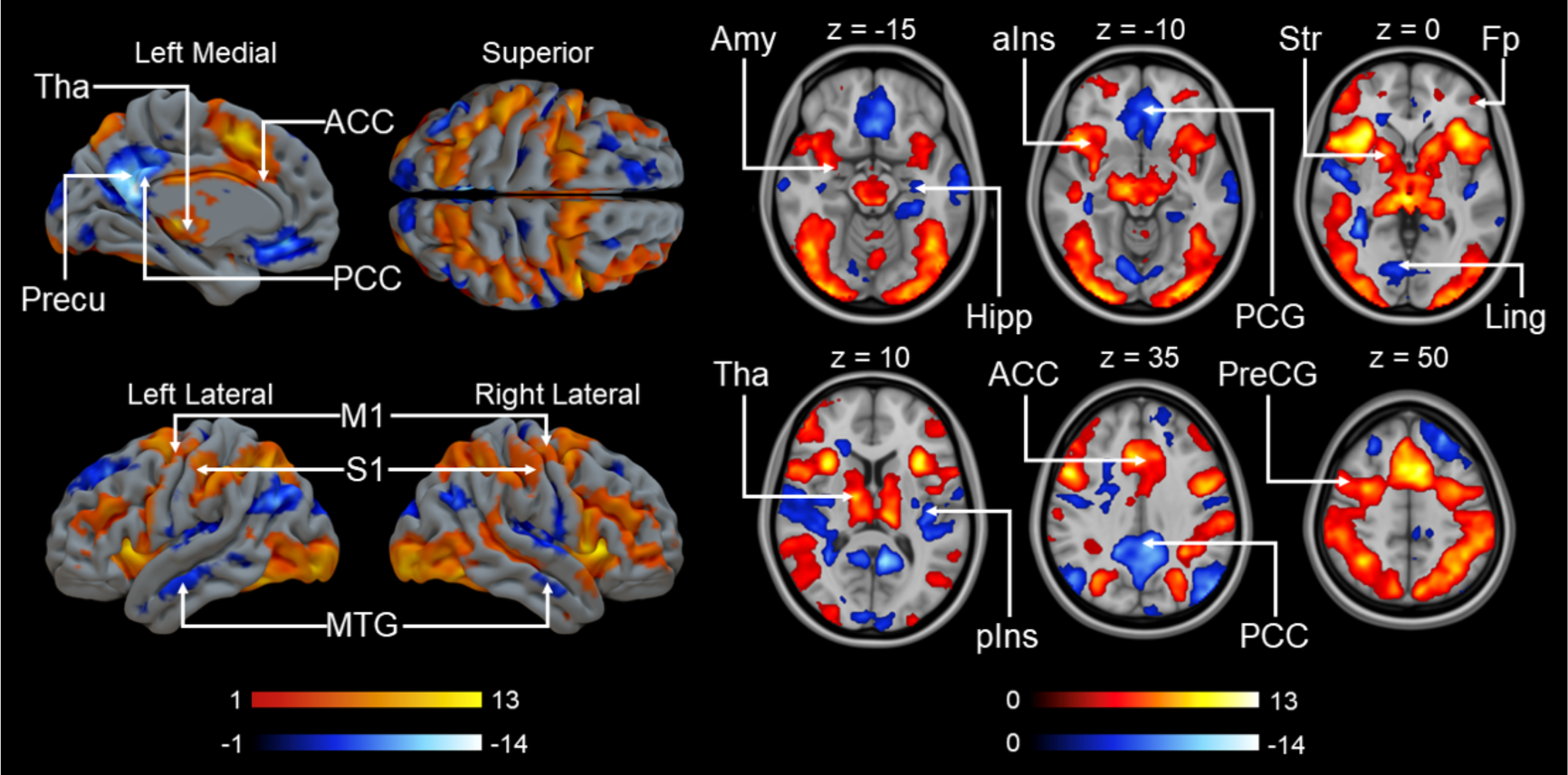
Main effect of brain responses to vicarious pain during the first video. The data are thresholded at p<0.001, FWE corrected at the cluster level (Positive: p_unc_<0.001, k=FWEc=114 voxels, 3.40<t<13.17; Negative: p_unc_<0.001, k=FWEc=97 voxels, 3.40<t<15.77). Colourbars indicate t statistic range. Tha = thalamus, Precu = precuneous cortex, ACC = anterior cingulate cortex, PCC = posterior cingulate cortex, M1 = primary motor cortex, S1 = primary somatosensory cortex, MTG = middle temporal gyrus, Amy = amygdala, Hipp = hippocampus, aIns = anterior insula, PCG = paracingulate gyrus, Str = striatum, Fp = frontal pole, Ling = lingual gyrus, pIns = posterior insula, PreCG = precentral gyrus.

**Figure 4.**
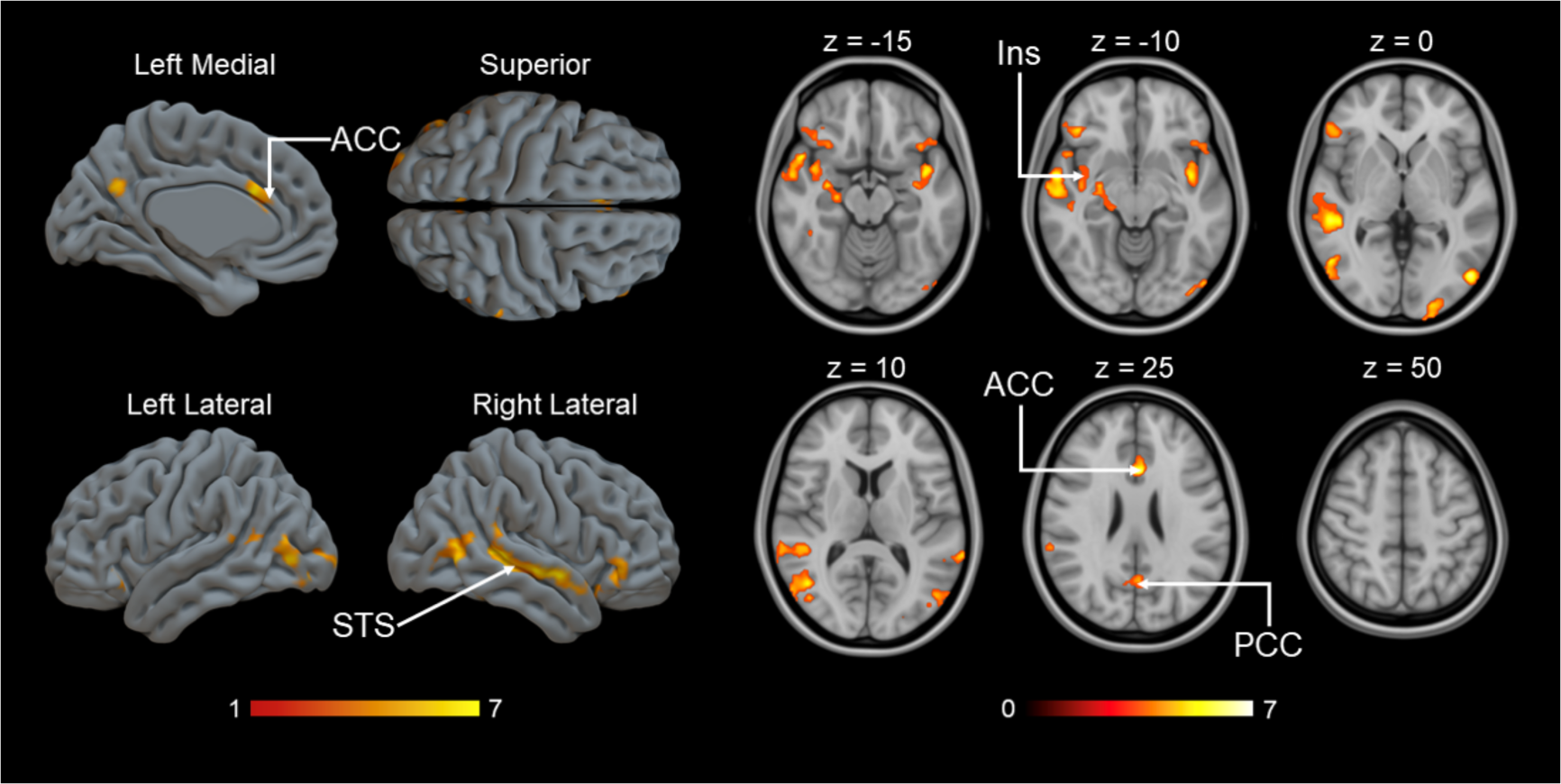
Brain regions with BOLD signals associated with donation size during the 1^st^ video. Colourbars indicate t statistic range. The data are thresholded at p<0.001, FWE corrected at the cluster level (p_unc_<0.001, k=FWEc=163 voxels, 3.40<t<7.20). ACC = anterior cingulate cortex, PCC = posterior cingulate cortex, STS = superior temporal sulcus, Ins = insula.

### BOLD-fMRI response changes following costly helping

In each trial, participants viewed two videos, one before their decision, and another following their decision. Comparing neural responses to the 1^st^ and 2^nd^ videos revealed decreased responses to the 2^nd^ video in areas involved in vicarious pain perception such as, insula, thalamus and ACC. Additional effects were found in striatum, M1 and S1 (**Fig. 5**). Because subjects were on average willing to help, the 2^nd^ video condition contained predominantly low shock intensity clips, thus direct comparison of 1^st^ and 2^nd^ could simply reflect differences in mean pain intensity across conditions. We subsequently restricted this analysis to low shock intensity levels (level 2 and level 3) in the 1^st^ video.

**Figure 5.**
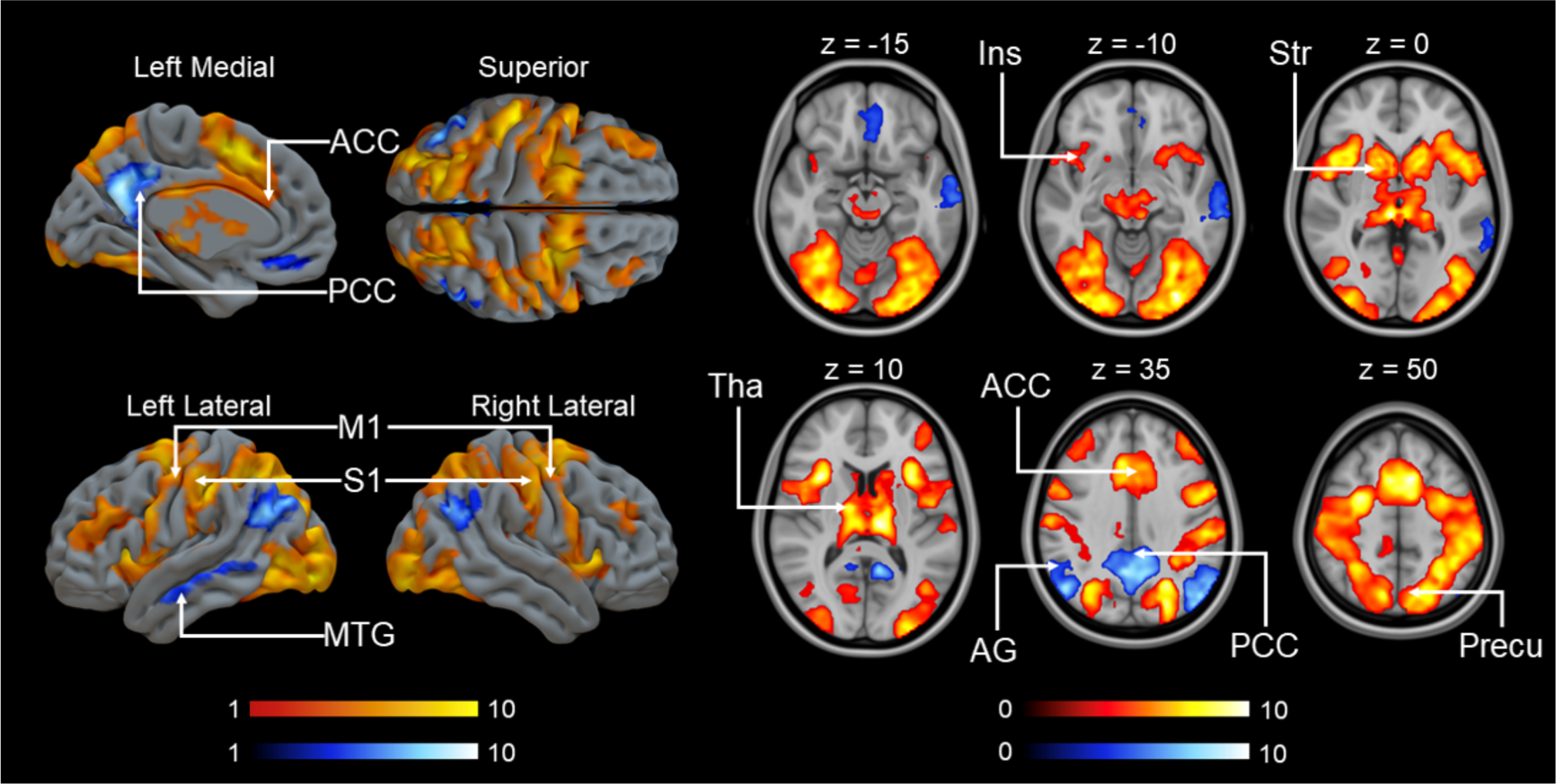
Brain activation for the 1^st^ versus the 2^nd^ video. The data are thresholded at p<0.001, FWE corrected at the cluster level (1^st^ > 2^nd^ : p_unc_<0.001, k=FWEc=692 voxels, 3.40<t<10.85; 2^nd^ > 1^st^ : p_unc_<0.001, k=FWEc=246 voxels, 3.40<t<10.24). Colourbars indicate t statistic range (red: 1^st^ > 2^nd^ video, blue: 1^st^ < 2^nd^ video). ACC = anterior cingulate cortex, PCC = posterior cingulate cortex, M1 = primary motor cortex, S1 = primary somatosensory cortex, MTG = middle temporal gyrus, Ins = insula, Str = striatum, Tha = thalamus, AG = angular gyrus, Precu = precuneus cortex.

Even when pain intensity was thus approximately matched across the 1^st^ and 2^nd^ videos (Fig. 6), effects were similar to those in Fig. 5, with thalamus, striatum, ACC, M1 and S1 showing decreased responses to the 2^nd^ video (**Fig. 6**).

**Figure 6.**
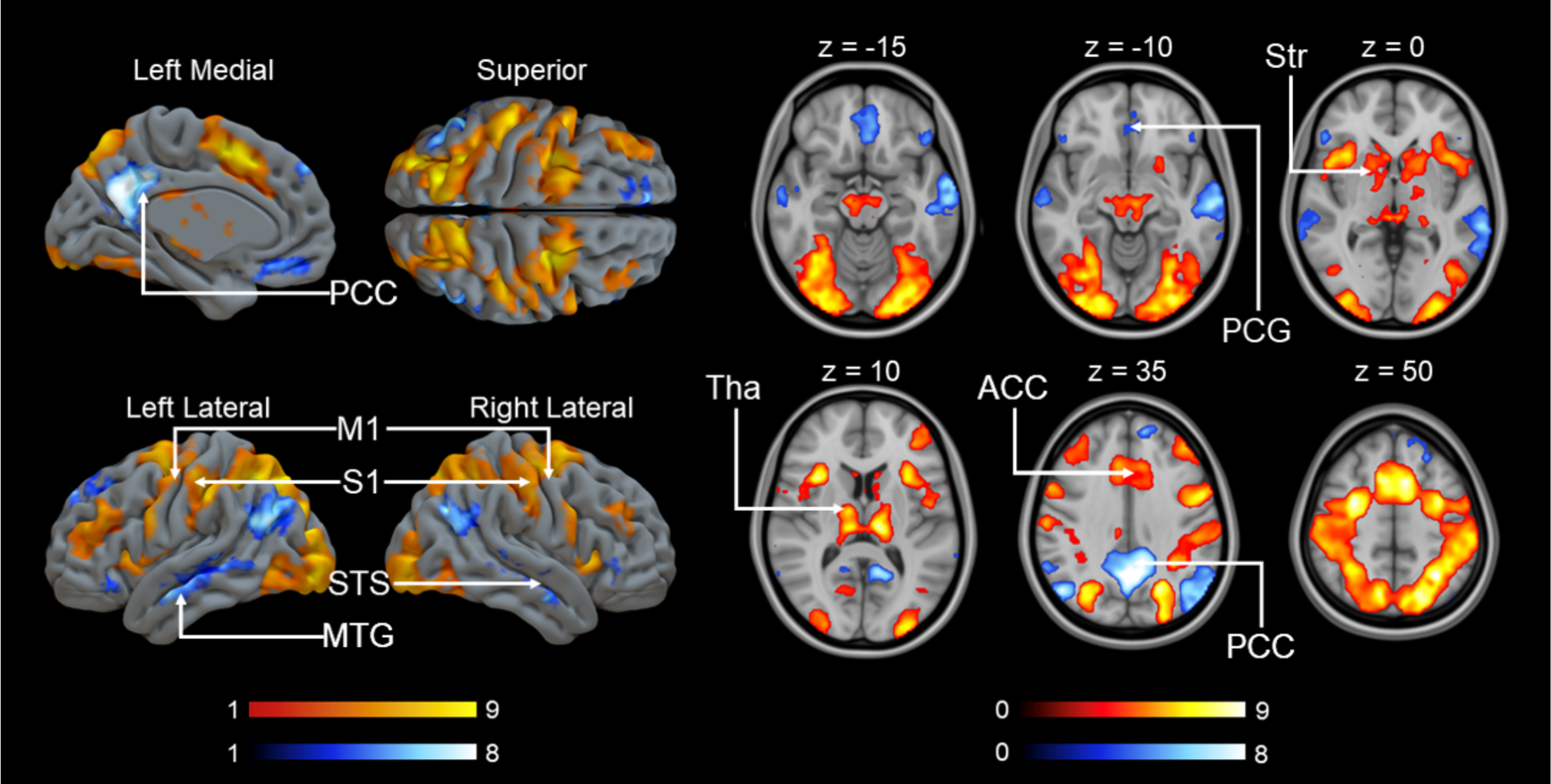
Brain activation for the low-level shock intensity trials in 1^st^ video versus the 2^nd^ video. The data are thresholded at p<0.001, FWE corrected at the cluster level (1^st^ > 2^nd^ : p_unc_<0.001, k=FWEc=146 voxels, 3.40<t<9.99; 2^nd^ > 1^st^ : p_unc_<0.001, k=FWEc=112 voxels, 3.40<t<10.40). Colourbars indicate t statistic range (red: 1^st^ > 2^nd^ video, blue: 1^st^ < 2^nd^ video). ACC = anterior cingulate cortex, PCC = posterior cingulate cortex, M1 = primary motor cortex, S1 = primary somatosensory cortex, STS = superior temporal sulcus, MTG = middle temporal gyrus, PCG = paracingulate gyrus, Str = striatum, Tha = thalamus.

### MOR-dependent responses to empathy for pain

To test our hypothesis that individual differences in MOR availability would be associated with differences in the neural circuitry associated with taking helping decisions, we next modelled BOLD responses associated with donation size during the 1^st^ video using regional MOR availabilities as predictors. We first used a PCA that captures the individual variability across the ROIs and extracted the first principal component that explained 60.7% of the individual variation in MOR binding (see **Table S1 for details**). The component scores were used as regressors to predict the haemodynamic response to donation size during the 1^st^ video. We found a generally negative correlation between BOLD signal and MOR availability, mainly in amygdala, striatum, insula, hippocampus, thalamus, anterior cingulate cortex and posterior cingulate cortex (**Fig. 7**). This indicates that participants with reduced MOR availability show BOLD signals in these regions that are more sensitive to trial-by-trial differences in donation. Importantly, these differences in the association between brain activity and donation as a function of MOR availability were observed despite similar donations across participants with higher or lower MOR availability (**Table 1**), ascertaining that these differences are not related to systematic differences in the predictors inserted into the first level fMRI models (see Lebreton et al., 2019 for a discussion of the importance of this factor).

**Figure 7.**
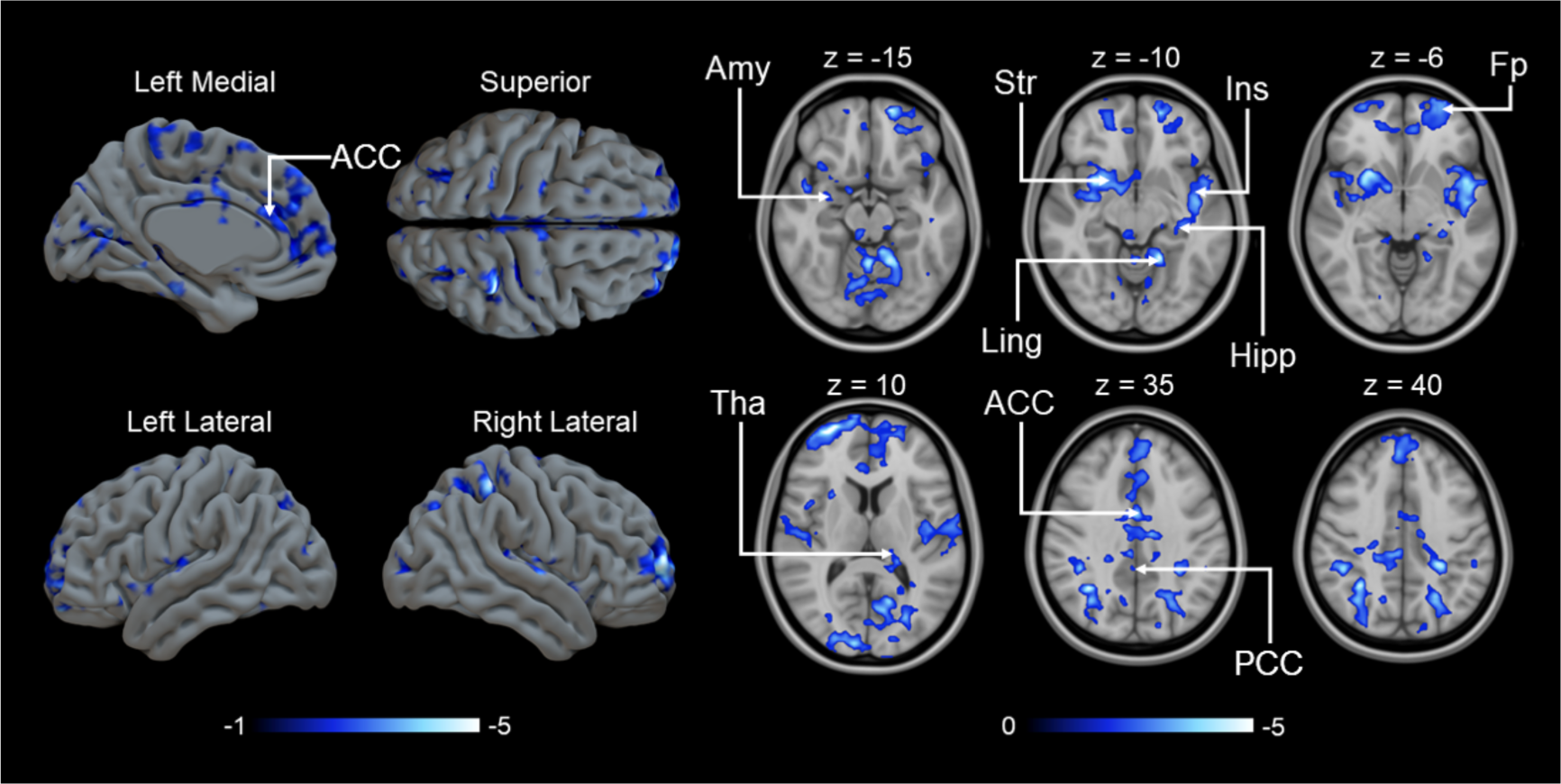
Negative correlation between the first component of MOR availability and the parametric modulator of BOLD response to donation size during 1^st^ video (p<0.05, FEW corrected at cluster level, p_unc_<0.05, k=FWEc=1800 voxels, 1.78<t<9.01). Amy = amygdala, Str = striatum, Ins = insula, Hipp = hippocampus, Ling = lingual gyrus, Fp = frontal pole, Tha = thalamus, ACC = anterior cingulate cortex, PCC = posterior cingulate cortex.

To focus more specifically on regions associated with witnessing the pain of others, we used the AVPS signature, which was found to be negatively associated with the third principal component (r = -0.659, p = 0.010, BF_10_=6.5) (**Fig. S5**). We also generated a cumulative map of correlation between regional MOR availabilities and BOLD responses to donation size in the 1^st^ video. The results replicated our prior study revealing a generally negative correlation between BOLD signal and MOR availability in vicarious pain related areas like the insula, anterior cingulate cortex and thalamus. Extensive association between MOR availability and brain activation was also observed in somatosensory areas, temporal gyrus, limbic regions and frontal cortex (**Fig. S2**) (Karjalainen et al., 2017).

### MOR-dependent responses to costly helping

Finally, we explored whether individual differences in MOR availability were associated with the BOLD activity during the donation phase, independently of what donation was given. As in the previous analysis, we used the 1^st^ PC scores for the MOR availability maps to predict haemodynamic activation during the donation phase. Unlike the negative associations between MOR and the parametric modulator for donation size during the 1^st^ video, this analysis revealed positive correlations between BOLD signal and MOR availability particularly in ACC and hippocampus (**Fig. 8**).

**Figure 8.**
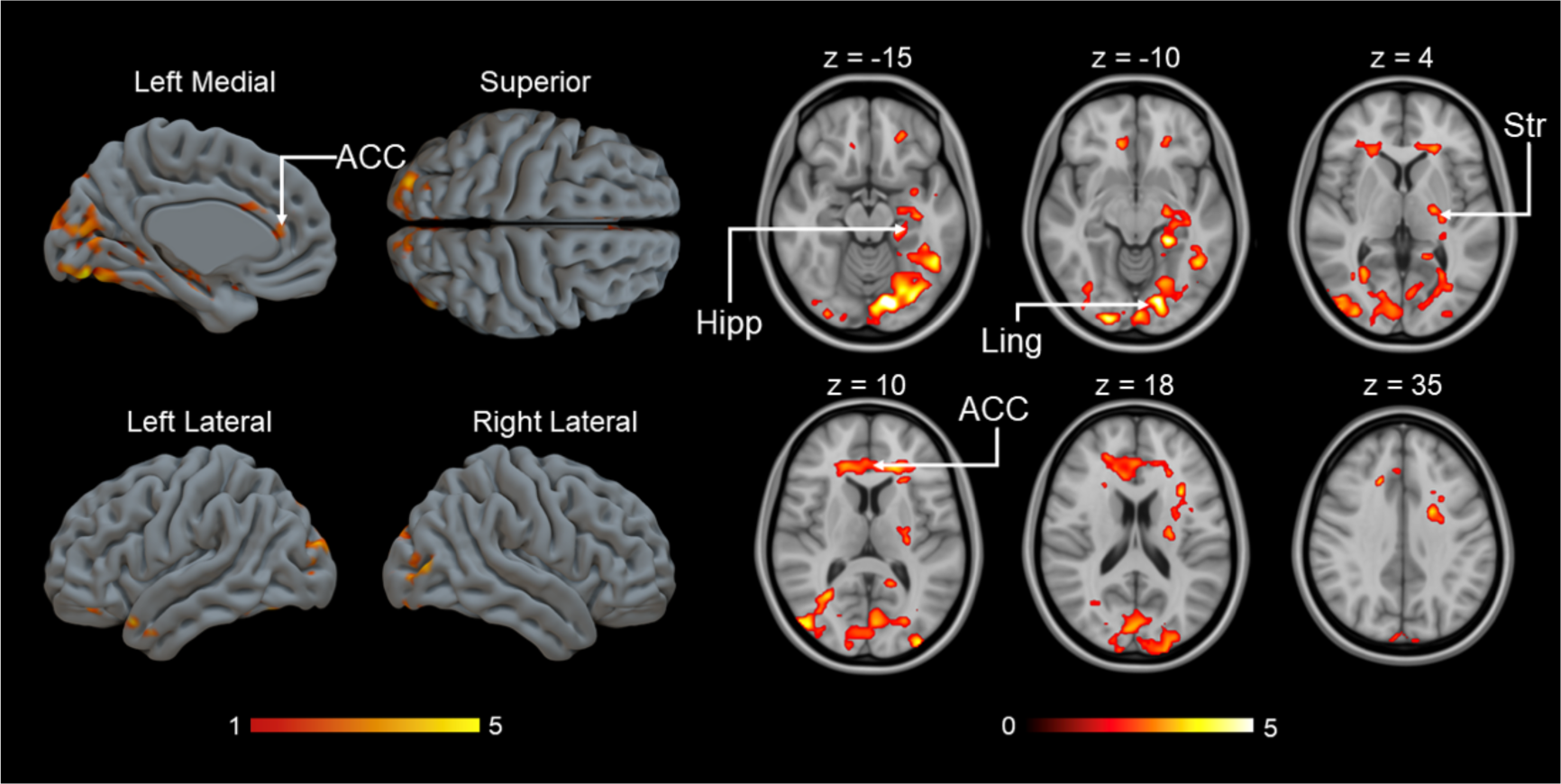
Positive correlation between MOR availability and BOLD response during the donation phase (p<0.05, FWE corrected at cluster level, Donation: p_unc_<0.05, k=FWEc=10316 voxels, 1.78<t<8.44). Hipp = hippocampus, ACC = anterior cingulate cortex, Ling = lingual gyrus, Str = striatum.

## Discussion

Our main finding was that individual differences in the endogenous opioid system tone do not directly alter participants’ decisions to help others, but are linked with brain activity differences during vicarious pain and costly, altruistic helping. Our subjects were generally willing to dive up significant amounts of money to help others and donated larger sums when they saw the confederate experiencing stronger pain. Trials in which participants donated more money were associated with increased BOLD activity in regions associated with empathy and the central nociceptive system (Ins, ACC, PCC STS). In line with studies showing that reduced baseline MOR availability is associated with heightened sensitivity to pain (Karjalainen et al., 2017), baseline MOR availability was negatively correlated with the BOLD responses while witnessing the pain of others in 1^st^ video associated with donating to help. Donation depended on how much pain was displayed in the 1^st^ video, and participants with reduced MOR availability had activity in circuits associated with empathy and nociception that were more strongly associated with donation. Also, brain activity during the donation phase, after they seeing how much pain the other participants expressed, depended on endogenous MOR tone in hippocampus, ACC, and striatum. These data provide the first direct *in vivo* evidence for the engagement of the opioid system in the neural processes occurring during costly helping, significantly extending the role of MORs in human social behavior.

### Neural basis of vicarious pain perception

As our goal was to model authentic altruistic helping that would also be costly to the participant, we used a naturalistic interactive setting in which we made participants believe that they were interacting with a real person whom they also met before the experiment. Behavioral data confirmed that the manipulation successfully induced costly helping: Participants donated more money when the shock intensity increased and the confederate expressed more intense pain. This finding accords with previous studies (Gallo et al., 2018) and suggests that people are willing to altruistically help strangers that they have met only recently. FMRI data revealed that witnessing the pain of the others during the 1^st^ video evoked widespread cortical and subcortical activation, in regions associated with empathic pain (aIns, ACC, M1, S1, thalamus, amygdala, and striatum). The results are consistent with meta-analyses of brain regions associated with empathy for pain (Jauniaux et al., 2019; Lamm et al., 2011; Timmers et al., 2018). ACC, PCC, insula, and STS response during 1^st^ video were stronger in trials in which participants later decided to donate more money. Within the limitations of fMRI, activity in these regions may thus have played a role in motivating costly helping. This dovetails with findings from previous studies that showed BOLD activity in similar networks of regions scaled with perceived pain in Hollywood-type videos with various intensities of pain (Karjalainen et al., 2017), and for the insula, recent intracranial recordings showing that the power in broadband gamma and the spiking of single neurons in this region scales with the perceived pain intensity for similar stimuli (Soyman et al., 2022). Importantly, this also matches findings using a similar task acquired in a different lab (Ioumpa et al., 2023).

### Endogenous opioids and vicarious pain perception

We observed a negative association between cerebral MOR availability and the relationship between BOLD activity and donation while viewing 1^st^ video across a wide range of brain regions involved in vicarious pain perception in three different ways. This association was significant in regions associated with empathy for pain (insula, ACC, thalamus, amygdala, and striatum). This negative association accords with previous findings associating lower MOR availability with higher sensitivity to pain (Hagelberg et al., 2012), more distress (Nummenmaa et al., 2020) as well as acute adverse emotions evoked by witnessing the pain of others (Karjalainen et al., 2017). Taken together these data suggest that lower MOR availability makes individuals more sensitive to suffer of others. These data also accord with the general role of the opioid system in maintaining social bonds and attachment (Manninen et al., 2017; Sun et al., 2022; Turtonen et al., 2021), which was here conceptualized as hemodynamic responses to seeing others in distress.

### Brain basis of costly helping

A peculiarity of the paradigm developed by Gallo et al. (2018) employed here is that participants can directly monitor the effectiveness of their costly helping by comparing 1^st^ video and 2^nd^ video. Comparing the brain activity across the 1^st^ and 2^nd^ videos (i.e., before and after helping), we found that the 1^st^ video evoked stronger activation in striatum, thalamus, ACC, M1, and S1, whether we compared all 1^st^ video against all 2^nd^ video, or repeat the analyses selecting instances in which 1^st^ video was of low intensity. This pattern was spatially similar to that elicited by the 1^st^ video only. One interpretation why the activity was lower in 2^nd^ video than 1^st^ video (even if the two videos showed similar levels of pain, Fig. 6) is that the opportunity to help decreased the vicarious pain response. This would accord with the negative state relief model for altruism, which states that helping others makes the helper feel better (Dovidio & Penner, 2003; Williamson & Clark, 1989). Altruistic behavior could thus be motivated by the (anticipated) alleviation of vicarious pain. Some have argued that altruism is driven by the rewarding nature of empathy and helping (Batson et al., 2014; Dovidio & Penner, 2003; FeldmanHall et al., 2015), which might predict helping-induced activations in the brain’s striatal reward systems (**Figure 5** and **supplementary figure S3**), which we, however, failed to find in our contrast between the 1^st^ and 2^nd^ videos.

Accordingly, within the limits of reverse inference, the neural data are perhaps best explained by the notion that helping is motivated by an anticipation of reduced vicarious pain/distress rather than by the anticipation of reward. It should, however, be noted that several alternative explanations could account for the observed reduction in brain responses from 1^st^ video to 2^nd^ video. First, 1^st^ video is relevant to the task given to the participants, and the association between the intensity of pain perceivable in 1^st^ video and the donation confirms that participants adapted their responses to the content of 1^st^ video. In contrast, 2^nd^ video is task-irrelevant. Task relevance and the attention it commands, may thus account for the intensity of the response to 1^st^ video. Second, the intensity of 1^st^ video is unpredictable, while the intensity of 2^nd^ video is predictable based on 1^st^ video and the donation. Given that a network similar to the one we observed here has been shown to encode prediction errors while witnessing the pain of others (Fornari et al., 2023), this difference in predictability may also account for the differences in BOLD responses. Together, the observed differences in BOLD activity across 1^st^ video and second should thus be interpreted with caution.

### Opioid system modulates brain activity during costly helping

As mentioned in the introduction, networks supporting costly helping have been described, but the role played by the opioid system remains poorly understood. Here we demonstrate that individual differences in baseline 𝜇-receptor availability also relate to the individual differences in brain activity while participants plan and report their costly-helping decision. We observed a predominantly positive correlation between cerebral MOR availability and an extensive BOLD response in the helping phase (donation) in hippocampus, ACC, and striatum. Prior PET studies have illustrated the modulatory role of the opioid system in emotion, vicarious pain, positive social interaction, prosocial motivation in humans, and prosocial behavior in monkeys (Fabre-Nys et al., 1982; Karjalainen et al., 2017; Keverne et al., 1989; Nummenmaa & Tuominen, 2018). Our findings thus broaden our knowledge of the function of the opioid system, demonstrating its modulating role in the neural processing of costly helping. Importantly, MOR availability had, in general, opposite relationship with haemodynamic responses in the cortical and subcortical pain and emotion circuits during vicarious pain perception (negative) and deciding to help (positive). This suggests that during vicarious pain perception, the MORs may act as a buffer against the stress evoked by seeing others in distress. Yet when relieving others’ stress via one’s own decisions becomes possible (here during donation phase), individuals with high MOR tone show amplified hemodynamic responses. Thus, individuals with high MOR tone might shift their brain activity from the moment of witnessing other people’s sufferance (Video1) to the moment of deciding to help them (decision phase). This suggests that MOR tone could be an important molecular pathway modulating altruism and sociability via multiple mechanisms.

Contrary to our findings that individual differences in brain activity during costly helping are associated with differences in MOR availability, we did not find a significant correlation between MOR availability and donation size. This is slightly unexpected given the association at the neural level with brain regions associated with donation size, but in general, brain-behaviour relationships require substantially larger samples than what can be achieved with invasive experimental PET studies (Marek et al., 2022). However, given the behavioural pharmacological studies pointing towards causal effects of opiates on helping (Hartmann et al., 2022) our data contributes to a nascent literature associating the MOR system with prosocial decision-making.

### Limitations

In a single PET scan, it is impossible to demonstrate the exact molecule-level mechanism for altered receptor availability (Kantonen, Karjalainen, Isoj, et al., 2020). Our single PET scan study design only allowed quantifying differences in baseline receptor binding, but not the capacity for endogenous opioid release. We also only studied females so the results may also not generalize to males, given sex differences in empathy (Christov-Moore et al., 2014; Hoffman, 1977) and MOR availability (Kantonen, Karjalainen, Isoj, et al., 2020). Finally, PET and fMRI data were not measured simultaneously, yet prior studies have established that [^11^C]carfentanil has excellent test-retest reliability even with multiple-months intervals.

## Conclusion

Placebo analgesia studies have suggested that the opioid system may contribute to costly helping decisions. We provide evidence that individual differences in 𝜇-opioid baseline availability can explain significant individual variability in how the brain processes the distress of others in a costly helping context, and how it processes the decision to help. Brain regions associated with empathic pain such as the anterior insula, ACC, thalamus, amygdala, striatum, were significantly activated while perceiving the pain of others, and more so on trials in which participants later decided to help more. Activation of these regions decreased following helping. MOR availability was negatively correlated with the processing of the pain of others but dominantly positively correlated with neural responses while making the decision to help. These results suggest that the opioid system is intimately involved in vicarious pain and neural processing of helping decisions.

## Acknowledgements

This work was supported by grants from the Academy of Finland (#332225), Sigrid Juselius Stiftelse, Signe och Ane Gyllenberg’s Stiftelse, and China Scholarship Council, as well as grants from the Dutch Research Council (NWO) to VG (VIDI grant 452-14-015) and CK (VICI grant 453-15-009).

## Supplementary Materials

### Stimuli

All videos played in the experiment (each video lasted 2s) were recorded in advance to span different shock levels (maximum to minimum: 6 to 2). All videos featured the same black background and black table. The participants could see the confederate’s face and right hand, and the intensity of the electric shock could only be detected by changes in the facial expression, mainly by change of eye brows and mouth. The location of the hand was kept similar cross movies using a mark on the table that was covered by the hand, and hence invisible to the viewers of the movies. When shooting the videos, rather than giving a real electric shock to the hand, we poked the actor on her thigh to induce her pain, as she reported this to be a more effective stimulus. Altogether, 17 of intensity 2, 12 of intensity 3, 9 of intensity 4, 7 of intensity 5 and 5 of intensity 6 were presented in the experiment randomly with fixed proportion in different intensity level. Pain levels were achieved by matching perceived pain levels to those used in Gallo et al. (2018) by author KI. Furthermore, there was significant positive relation between shock intensity in 1^st^ video and donation indicating the stimuli worked well (r = 0.672, *p* < 0.001). For 1^st^ video, the number of different shock intensity condition were similar, and reduced high shock intensity video in 2^nd^ video caused by participants’ donation, see **Figure S1**.

**Figure S1.**
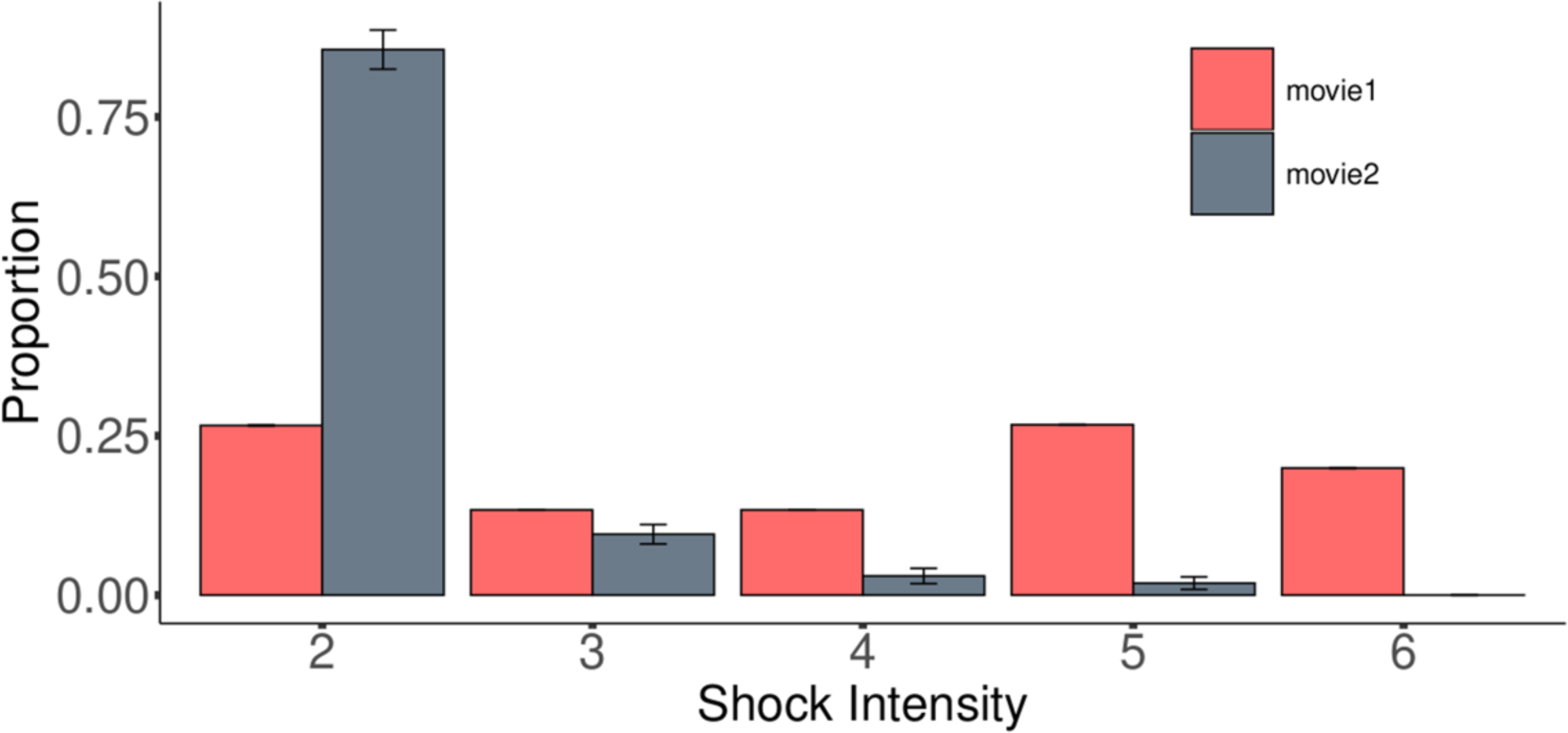
The proportion of shock intensity for 1^st^ video and 2^nd^ video. Error bar represents *SE*.

### MOR-dependent responses to pain

Instead of using principal component analysis, we directly cumulate all the significate ROIs after corrected (**Fig. S2**).

**Figure S2.**
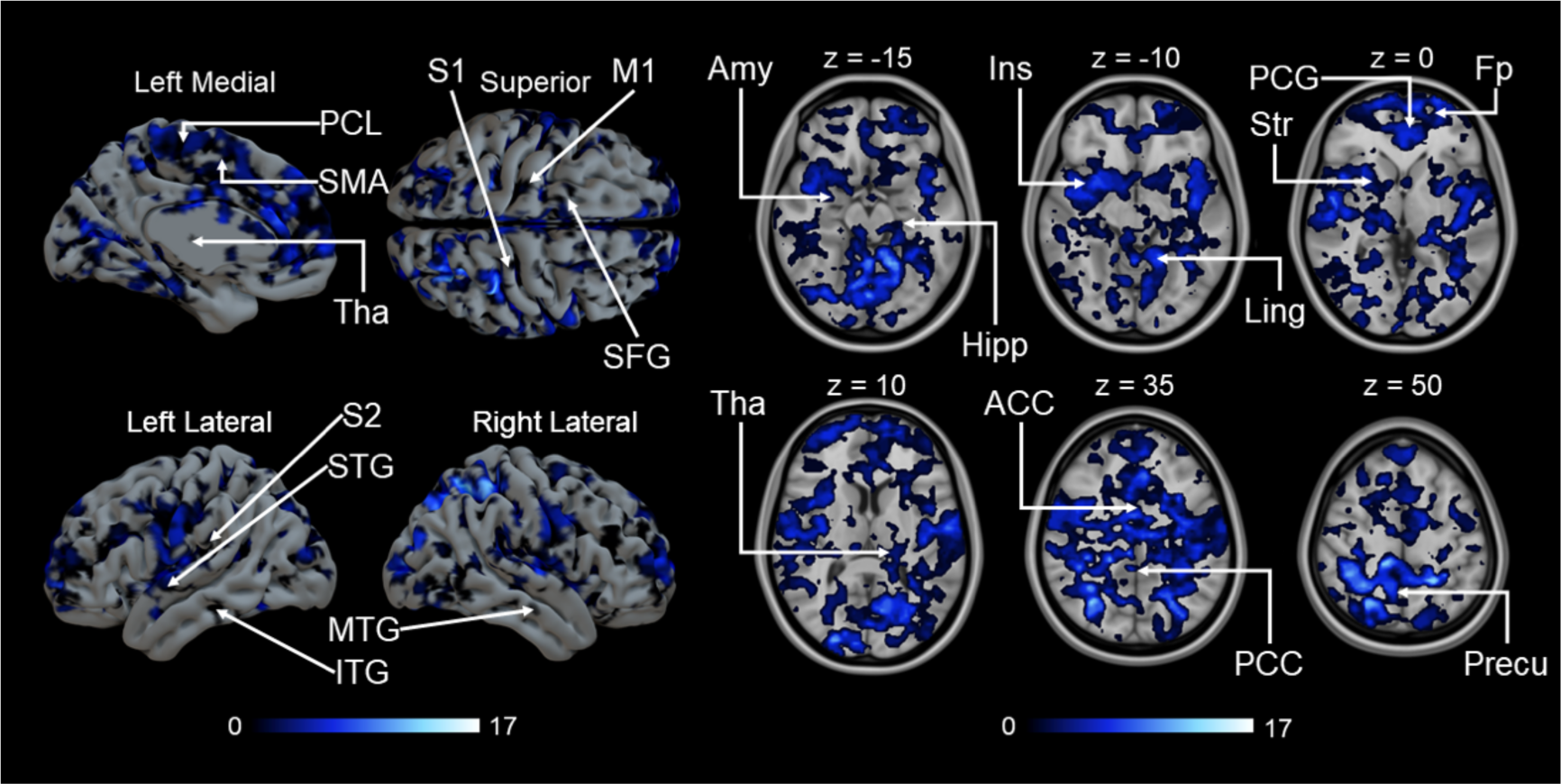
Cumulative maps showing the number of ROIs (out of 17) whose [^11^C]carfentanil *BP*_ND_ was negatively correlated (p<0.05, FWE corrected at cluster level) with BOLD response to donation size. PCL = paracentral lobule, SMA = supplementary motor area, Tha = thalamus, S1 = primary somatosensory cortex, M1 = primary motor cortex, SFG = superior frontal gyrus, S2 = secondary somatosensory cortex, STG = superior temporal gyrus, MTG = middle temporal gyrus, ITG = inferior temporal gyrus, Amy = amygdala, Hipp = hippocampus, Ins = insula, Ling = lingual gyrus, PCG = paracingulate gyrus, Str = striatum, Fp = frontal pole, ACC = anterior cingulate cortex, PCC = posterior cingulate cortex, Precu = precuneous cortex.

### Vicarious pain responses to 2^nd^ video

After donation, BOLD response to 2^nd^ video was different from 1^st^ video. Anterior insula, and angular gyrus were activated. The anterior insula in particular has been associated with vicarious pain activated in previous studies (**Fig. S3**).

**Figure S3.**
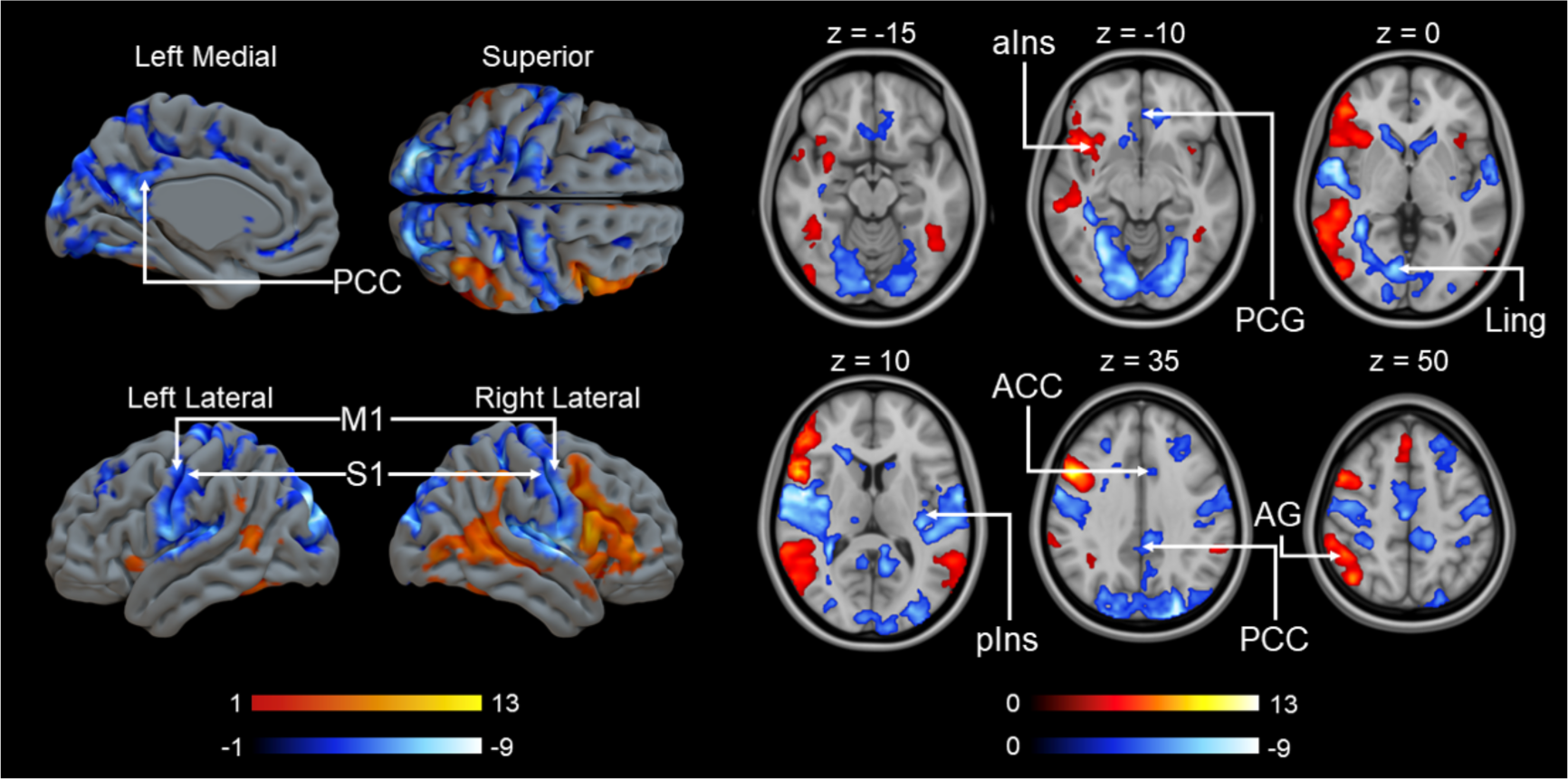
Main effect of pain perception in 2^nd^ video. Colourbars indicate t statistic range. The data are thresholded at p<0.001, FWE corrected at the cluster level (Positive: p_unc_<0.001, k=FWEc=110 voxels, 3.40<t<12.89; Negative: p_unc_<0.001, k=FWEc=121 voxels, 3.40<t<10.62). PCC = posterior cingulate cortex, M1 = primary motor cortex, S1 = primary somatosensory cortex, aIns = anterior insula, PCG = paracingulate gyrus, Ling = lingual gyrus, pIns = posterior insula, ACC = anterior cingulate cortex, AG = angular gyrus.

### Brain responses in donation phase

In donation phase, as shown in **Figure S4**. Extensive brain areas were activated, including anterior insula, striatum, thalamus, anterior cingulate cortex, M1, S1.

**Figure S4.**
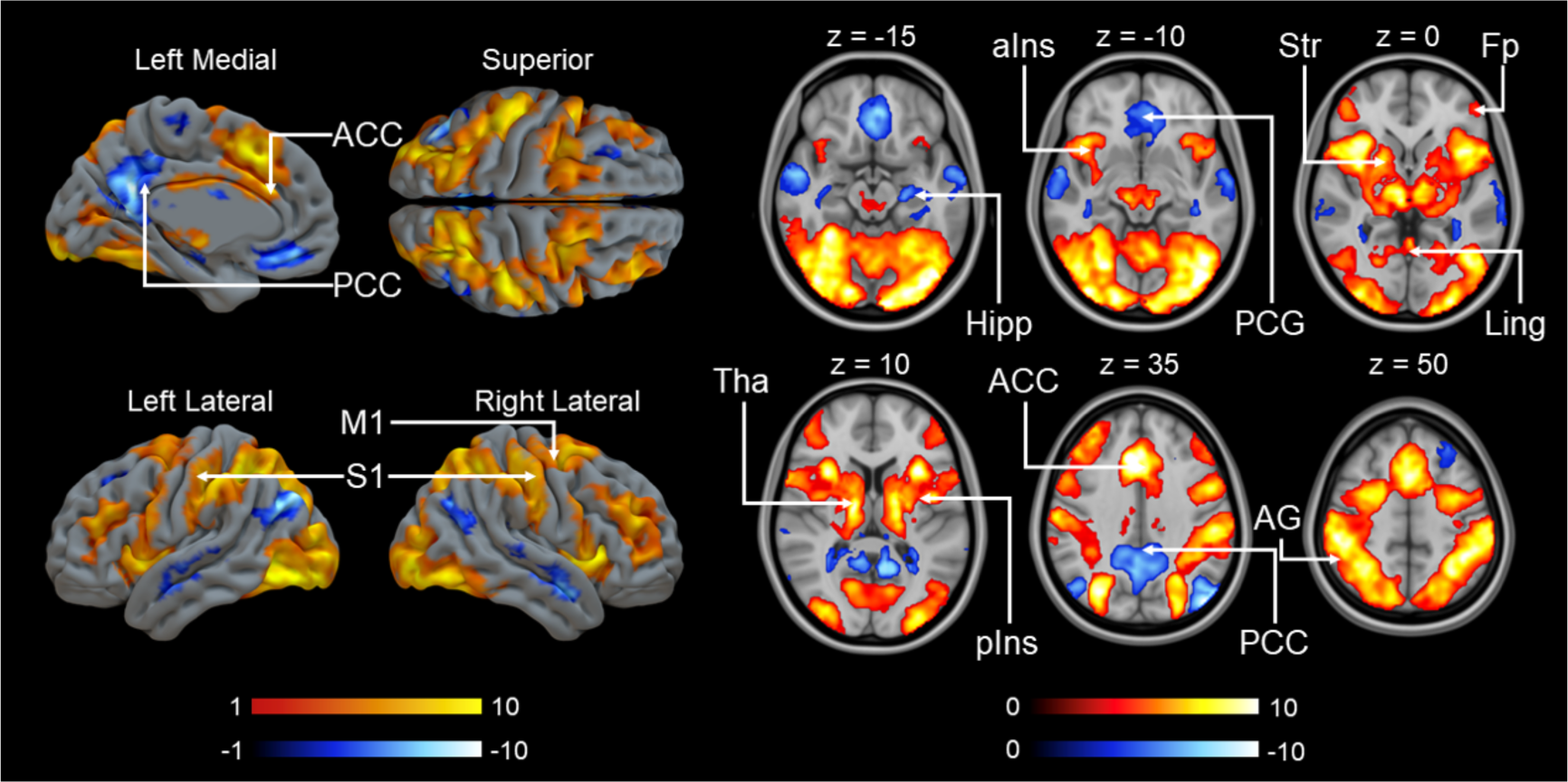
Main effect of brain activation to donation phase. Colourbars indicate t statistic range. The data are thresholded at p<0.001, FWE corrected at the cluster level (Positive: p_unc_<0.001, k=FWEc=67990 voxels, 3.40<t<14.73; Negative: p_unc_<0.001, k=FWEc=221 voxels, 3.40<t<11.12). ACC = anterior cingulate cortex, PCC = posterior cingulate cortex, M1 = primary motor cortex, S1 = primary somatosensory cortex, Hipp = hippocampus, aIns = anterior insula, PCG = paracingulate gyrus, Str = striatum, Fp = frontal pole, Ling = lingual gyrus, Tha = thalamus, pIns = posterior, AG = angular gyrus.

### Principal component analysis results

PCA result shows that the first three component can explain 90.1% of the variance, from the first component to the third component, they respectively explained 60.74%, 22.13%, and 7.27% of the variance. See the loadings of each ROIs below.

**Table S1.** Loadings of the first three principal component on the ROIs. Amy = amygdala, cau = caudate, cer = cerebellum, dacc = dorsal anterior cingulate cortex, inftemp = inferior temporal gyrus, ins = insula, midtempo = middle temporal gyrus, nacc = nucleus accumbens, ofc = orbitofrontal cortex, parsop = pars opercularis, pcc = posterior cingulate cortex, put = putamen, racc = rostral anterior cingulate cortex, supfront = superior frontal gyrus, suptemp = superior temporal gyrus, tempol = temporal pole, tha = thalamus.

| ROI | PC1 | PC2 | PC3 |
| --- | --- | --- | --- |
| amy | 0.317 | 0.632 | -0.207 |
| cau | 0.149 | 0.121 | -0.386 |
| cer | 0.231 | 0.254 | -0.069 |
| dacc | 0.312 | 0.004 | -0.024 |
| inftemp | 0.219 | -0.010 | 0.152 |
| ins | 0.214 | 0.212 | -0.091 |
| midtemp | 0.200 | -0.042 | 0.091 |
| nacc | 0.547 | 0.040 | 0.607 |
| ofc | 0.248 | -0.385 | 0.184 |
| parsop | 0.154 | -0.218 | -0.084 |
| pcc | 0.169 | -0.224 | -0.091 |
| put | 0.110 | -0.248 | -0.276 |
| racc | 0.230 | -0.248 | -0.178 |
| supfront | 0.120 | -0.171 | -0.210 |
| suptemp | 0.121 | -0.188 | -0.131 |
| tempol | 0.168 | -0.197 | -0.224 |
| tha | 0.239 | -0.030 | -0.355 |

**Figure S5.**
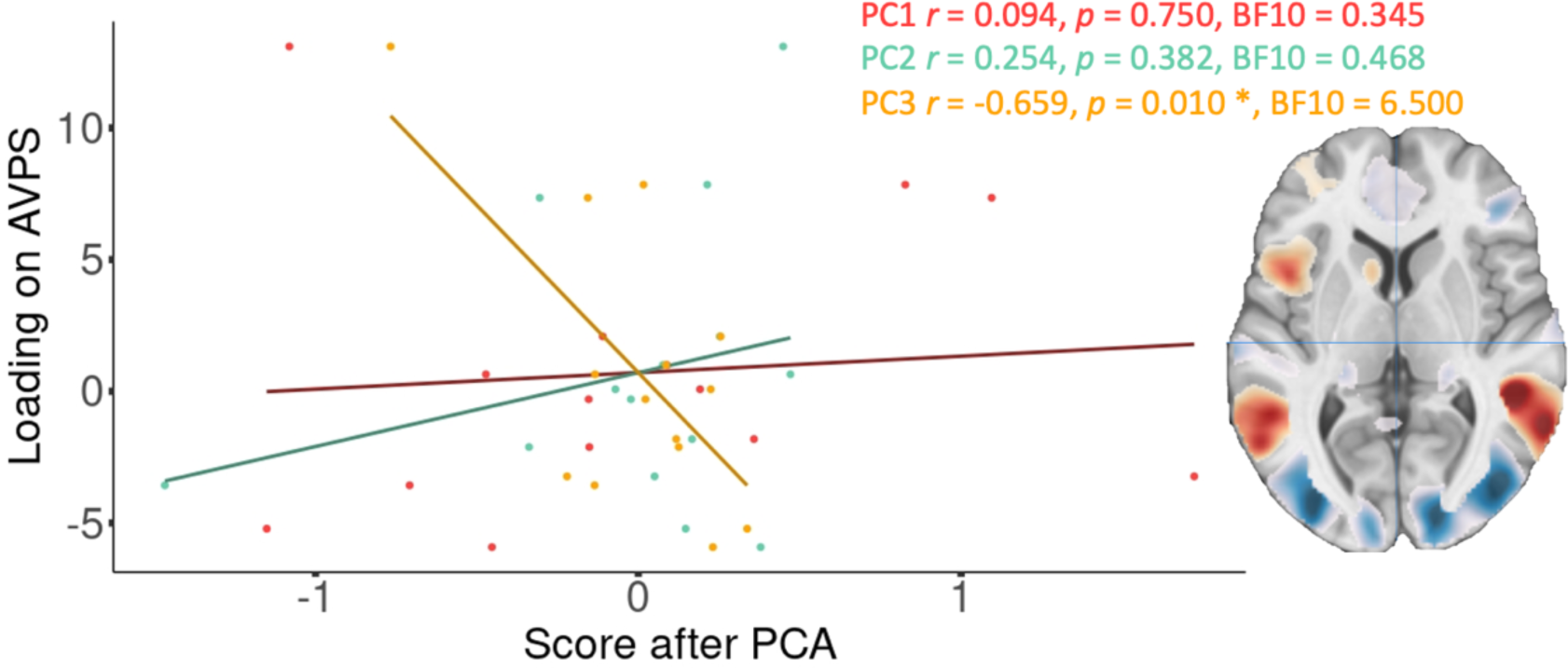
Correlation between the loading on AVPS of the parametric modulator donation size during 1^st^ video and first three principal components. (Red=PC1, green=PC2, yellow=PC3). The sagittal slice on the right represents a thresholded version of the AVPS signature (https://identifiers.org/neurovault.collection:6332) that was dot-multiplied with each participants parametric modulator volume. Each dot represents the loading of one participant onto the AVPS.

